# Phylogenetic relationships of the Helmeted Woodpecker (*Dryocopus galeatus*): A case of interspecific mimicry?

**DOI:** 10.1101/023663

**Authors:** Brett W. Benz, Mark B. Robbins, Kevin J. Zimmer

## Abstract

Examples of phenotypic convergence in plumage coloration have been reported in a wide diversity of avian taxonomic groups, yet the underlying evolutionary mechanisms driving this phenomenon have received little scientific inquiry. Herein, we document a striking new case of plumage convergence in the Helmeted Woodpecker (*Dryocopus galeatus*) and explore the possibility of visual mimicry among Atlantic Forest woodpeckers. Our multi-locus phylogenetic analyses unequivocally place *D*. *galeatus* within *Celeus*, indicating the former has subsequently converged in appearance upon the distantly related and syntopic *Dryocopus lineatus*, to which it bears a remarkable resemblance in plumage coloration and pattern. Although details of the Helmeted Woodpecker’s ecology and natural history are only now beginning to emerge, its smaller size and submissive behavior are consistent with predictions derived from evolutionary game theory models and the interspecific social dominance mimicry hypothesis (ISDM). Moreover, estimates of avian visual acuity suggest that size-related mimetic deception is plausible at distances ecologically relevant to *Celeus* and *Dryocopus* foraging behavior. In light of our results, we recommend taxonomic transfer of *D*. *galeatus* to *Celeus* and emphasize the need for detailed behavioral studies that examine the social costs and benefits of plumage convergence to explicitly test for ISDM and other forms of mimicry in these Atlantic Forest woodpecker communities. Future field studies examining potential cases of competitive mimicry should also take into account the mimic’s acoustic behavior, particularly in the presence of putative model species and other heterospecific competitors, as any discontinuity between morphological and behavioral mimicry would likely preclude the possibility of deception.

## INTRODUCTION

Recent progress in assembling the avian tree of life has shed light on numerous instances of non-aposematic plumage convergence in disparate taxonomic groups (Weibel and Moore 2005, Weckstein 2005, Tello et al. 2009, Jønsson et al. 2010). The highest incidence and perhaps most comprehensive examples of this phenomenon occur within the woodpeckers (Picidae), which exhibit convergent evolution of elaborate plumage patterns in at least eleven genera (Webb and Moore 2005, Weibel and Moore 2005, Benz et al. 2006, Moore et al. 2006, Fuchs et al. 2008). Although interspecific visual mimicry has long been suspected in various cases of avian plumage convergence, the adaptive significance and underlying socio-ecological mechanisms that promote phenotypic similarity in the absence of aposematism have received little attention and remain poorly understood by comparison with other forms of mimicry (Wallace 1869, Cody 1969, Diamond 1982, Ruxton et al. 2004, Rainey and Grether 2007).

Alfred Russell Wallace (1869) hypothesized that visual mimicry may explain the serial evolution of plumage convergence between Old World orioles (*Oriolus*) and Australasian friarbirds (*Philemon*) co-distributed across Wallacea. Assuming a classic three-player system comprised of a model, mimic, and third party observer, Wallace reasoned that the smaller, subordinate *Oriolus* species were mimicking the appearance of larger, highly aggressive friarbirds to avoid attack from hawks or other socially dominant non-model species. An alternative hypothesis was proposed over a century later by Jared Diamond (1982) who argued that the subordinate *Oriolus* mimics were instead incurring social benefits by directly deceiving the *Philemon* models, thereby minimizing interspecific aggression at highly contested foraging sites and gaining access to nectar resources that would otherwise be unavailable. Diamond’s two player hypothesis was largely based on cursory field observations of social interactions between *Philemon* and *Oriolus* species across the Australo-Papuan region, thus he remained unclear whether third party deception was also necessary to promote and maintain competitive mimicry.

Other researchers have invoked hypotheses of natural selection for enhanced interspecific signaling to explain convergent evolution of phenotypic similarities in birds. Martin Moynihan (1968) theorized that convergence in plumage coloration may foster more efficient interspecific communication within mixed-species foraging flocks by co-opting adaptive signal-receiver biases. By contrast, Martin Cody (1969) proposed that phenotypic similarities may actually enhance interspecific territoriality between ecological competitors by eliciting heightened aggression. He examined several cases of plumage convergence within woodpeckers (*Dinopium*-*Chrysocolaptes*, *Meiglyptes*-*Hemicircus*, *Dryocopus*-*Campephilus, Micropternus*-*Blythipicus*) as well as African bush-shrikes (*Chlorophoneus*-*Malaconotus*), and reasoned that visual mimicry would promote more efficient interspecific communication and exclusion of potential ecological competitors given these same social signals presumably form the basis for conspecific territorial interactions. However, this hypothesis has received criticism for its inconsistency with competitive exclusion theory and failure to distinguish when convergent evolution should be favored over character displacement (Murray 1976, Prum 2014).

The primary theoretical deficiencies inherent to these previous works have recently been addressed by Prum and Samuelson (2012) who developed an explicit evolutionary framework derived from game theory to examine the selection forces associated with non-aposematic visual mimicry. Elaborating upon the classic hawk-dove game, they used a well-documented case of plumage convergence between two North American woodpeckers (*Picoides villosus* and *Picoides pubescens*; Weibel and Moore 2005) to estimate the coevolutionary fitness dynamics between model and mimic, thereby establishing the conditions that promote evolution of interspecific social dominance mimicry (ISDM). Prum and Samuelson (2012) define ISDM as a type of social parasitism in which a smaller subordinate species uses visual deception to minimize competitive interference with a dominant model taxon and gain access to enhanced feeding opportunities. Several predictions with respect to the players’ ecology, behavior, and body size have emerged from these analyses that should further facilitate testing ISDM in birds and other vertebrate groups. First, mimetic species are smaller and socially subordinate to their model counterparts, yet these size differences are constrained such that visual deception must be feasible at distances germane to the players’ behavioral ecology. Second, the costs of mimicry and value of contested resources cannot be exceedingly high for mimic and model species to coexist through time. Third, shared similarities in appearance are not attributed to homologous traits, in that the mimic and model are not closely related sister species. Lastly, both mimic and model are under natural selection to maintain or evade visual deception, respectively. As such, coevolutionary radiations may emerge if the evolution of mimicry precedes diversification in the model species. In the present study, we examine the evidence for ISDM and alternative explanations of phenotypic convergence in the Helmeted Woodpecker (*Dryocopus galeatus*), a little known Atlantic Forest endemic whose systematic affinities remain unclear given its enigmatic combination of morphological and behavioral characters.

Initially described as *Picus galeatus* (Temminck 1822), the Helmeted Woodpecker was soon transferred to *Dryocopus* by Gray (1845) where it has generally remained, but not without comment. Short (1982) was apparently the first to recognize that *D*. *galeatus* shares morphological characters both with *Dryocopus* and *Celeus*, commenting that the species is “beautifully intermediate” and could be placed in either genus. Specifically, he notes the weaker curved bill, exposed nostrils, cinnamon wing linings, and white upper tail coverts which are characteristic of *Celeus*, whereas the uniform dark dorsal plumage, ventral barring, whitish neck stripe, unbarred wings, and fully red crest are traits shared with Neotropical *Dryocopus*. Despite these morphological similarities to *Celeus*, Short concluded that *D*. *galeatus* is most likely sister to the *D*. *schulzi* + *D*. *lineatus* clade, to which it bears strong phenotypic resemblance (Figure 1) and is narrowly syntopic (Figure 2) with the latter taxon (Short 1982, Winkler et al. 1995). Herein, we build upon a recent molecular phylogenetic analysis of *Celeus* by incorporating multi locus sequence data from *D*. *galeatus* and putative congenerics to resolve its systematic position and provide a phylogenetic basis for examining plumage evolution.

**FIGURE 1.**
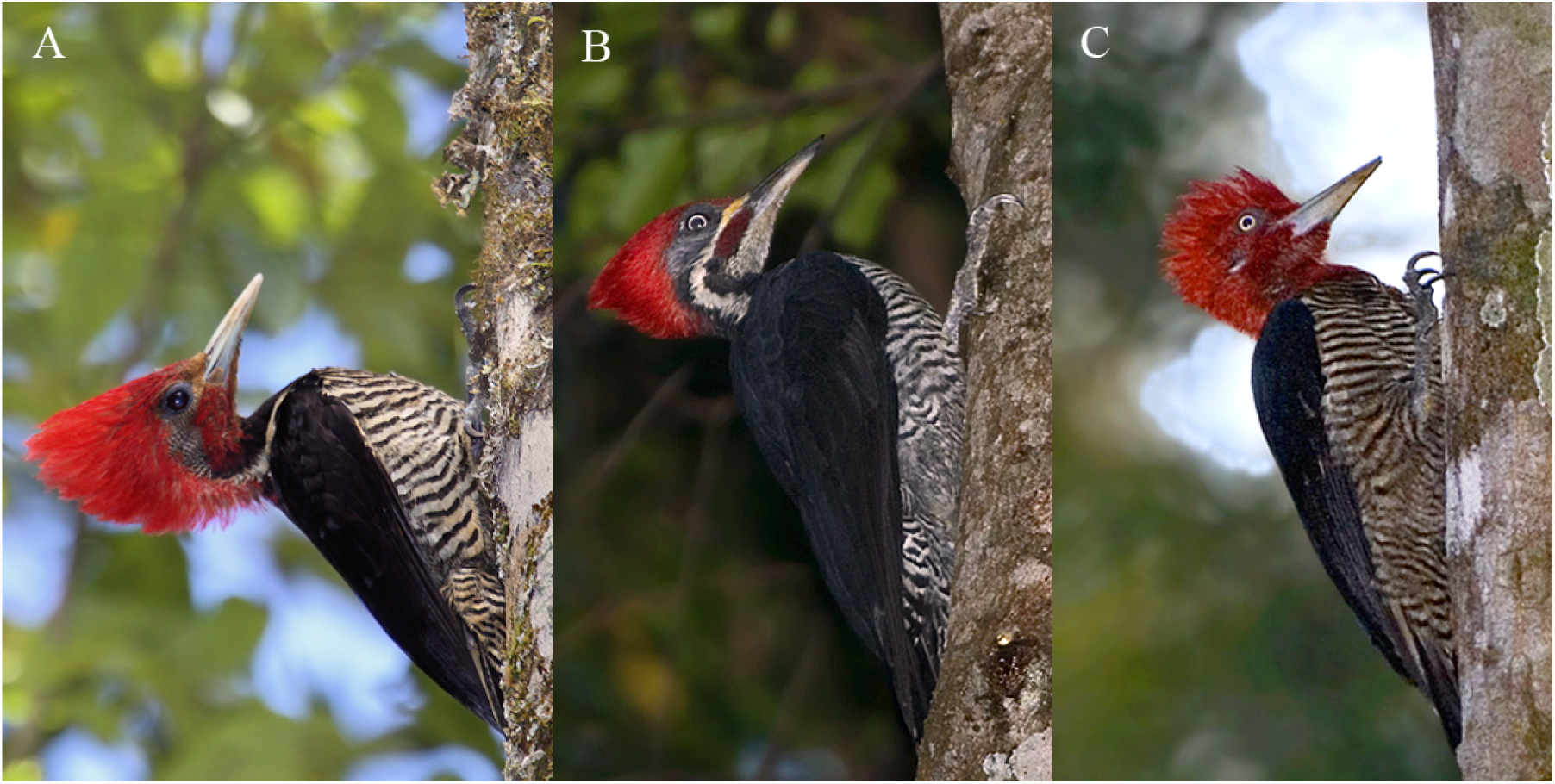
Images of adult male (A) *Dryocopus galeatus*, (B) *Dryocopus lineatus erythrops*, and (C) *Campephilus robustu*s. Although highly similar in appearance, *D*. *galeatus* weighs less than half of either *D*. *lineatus* or *C*. *robustus* and is 22-25% smaller in length respectively. Photographs are reproduced with permission from K. J. Zimmer [(A) taken in Intervales State Park, São Paulo, Brazil] and Ricardo Jensen Moller [(B, C) photographed in Misiones, Argentina].

**FIGURE 2.**
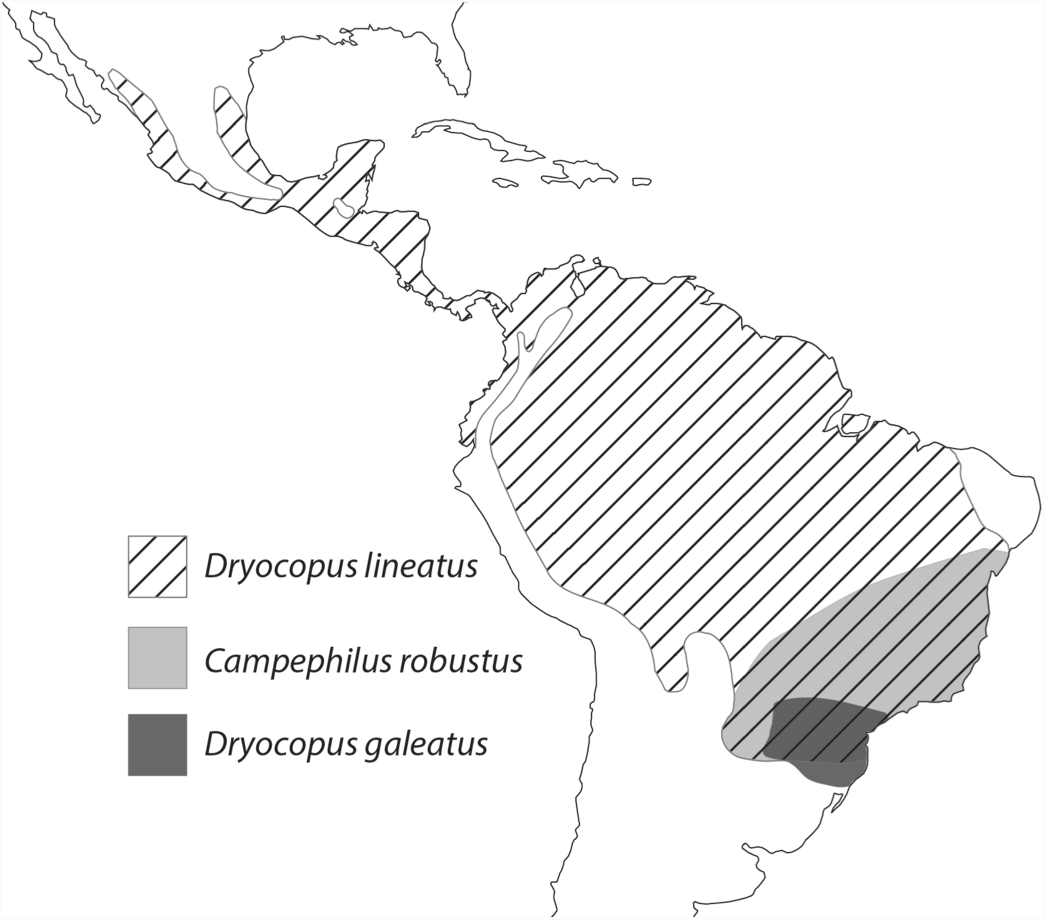
Approximate distributions of *Dryocopus galeatus*, *Dryocopus lineatus*, and *Campephilus robustus*, illustrating the potential for sympatry and social interactions between these taxa.

## METHODS

### Taxon Sampling and Sequencing

We obtained molecular sequence data from 44 woodpecker specimens representing 32 species in 14 genera (Appendix A). Taxon sampling was concentrated within the Malarpicini, including 5 of 7 *Dryocopus* species, all currently recognized members of *Celeus* (Benz and Robbins 2011), and at least one species from each of the remaining Neotropical genera (Winkler and Christie 2002). Six outgroup taxa were selected from the Megapicini and Dendropicini based on previous molecular phylogenetic studies of the Picidae (Webb and Moore 2005, Benz et al. 2006, Fuchs et al. 2007). Whole genomic DNA was extracted from muscle tissue using proteinase K digestion under manufacturer’s protocols (DNeasy tissue kit, Qiagen). We employed standard PCR amplification and Sanger sequencing methods to generate sequence data from four mitochondrial genes (NADH dehydrogenase subunits 2 and 3 [ND2 1041 bp; ND3 351 bp], ATP synthase subunits 6 and 8 [ATP6, 684 bp; ATP8, 168 bp]) and two nuclear loci (intron 7 of the β-fibrinogen gene [β-FIBI7, 913 bp], and a segment of the nonhistone chromosomal protein HMG-17 gene including exon 2 and adjacent mRNAs [HMGN2, 709 bp]). Sequence data was obtained from ND2, ND3, and HMGN2 for all 44 specimens, whereas sequencing effort for β-FIBI7 was limited to contemporary samples (n=41) and that of ATP6-8 to members of *Celeus* and *Dryocopus* (n=27). Ancient DNA sequencing techniques were used to obtain complete ND2, ND3, ATP6-8 and HMGN2 sequence data from two museum specimens of *D*. *galeatus* collected in 1959 from Tobunas, Argentina (Appendix A). See Benz and Robbins (2011) for comprehensive details of the laboratory protocols employed herein. GenBank accession numbers for all sequence data generated prior to this investigation can be found in the aforementioned 2011 study and Benz et al. (2006).

### Phylogenetic analysis

We used the Akaike Information Criteria (AIC) implemented in jMODELTEST 2 (Guindon and Gascuel 2003, Darriba et al. 2012) to determine best-fit models of evolution for individual nuclear loci and the concatenated mtDNA data set partitioned by codon position. A series of Bayesian phylogenetic analyses were conducted for each of these data sets in MRBAYES 3.2.1 (Ronquist et al. 2012) to assess potential conflict in phylogenetic signal among individual gene trees. We used a flat default prior distribution for parameter estimation and mitochondrial data partitions were permitted to vary independently by unlinking all parameters except topology and branch length. Two independent analyses were run per locus for 2 × 10^7^ generations and sampled every 100 generations, resulting in a total of 2 × 10^5^ samples. Stationarity for each analysis was assessed by examining average standard deviation of split frequencies, plotting model parameter posterior probability densities in TRACER v. 1.5 (Rambaut and Drummond 2007), and examining clade posterior probabilities across runs using the compare and slide functions in AWTY (Nylander et al. 2008). Trees that were sampled prior to the analysis reaching stationarity were discarded as a burnin. These same run parameters were then used for a combined data analysis with sequences partitioned by nuclear loci and mitochondrial codon position.

Maximum likelihood analyses of the individual loci and five-partition data matrix were conducted in GARLI v2.0 (Zwickl 2006) to provide alternative estimates of topology and node support. A total of thirty runs were conducted under default parameters to ensure the optimal –lnL solution had been reached, and topologies were selected after 10,000 generations with no significant improvement in –lnL (improvement values set at 0.01 with a total improvement lower than 0.05 compared to the last topology recovered). Node support was assessed using 500 non-parametric bootstrap replicates that were run with the above default parameters.

### Plumage analysis

We examined multiple museum specimens of *D*. *galeatus* (n=8) and candidate model species *D*. *lineatus* (n=14) and *C*. *robustus* (n=5) to assess overall plumage similarities and the potential for visual deception between these codistributed taxa. Representatives of each of the five recognized *D*. *lineatus* subspecies were examined, including four individuals of *D*. *lineatus erythrops*, the southern most taxon in the *lineatus* complex and the only subspecies co-distributed with *D*. *galeatus*. To examine the distribution of key plumage traits more broadly within the Malarpicini, we examined multiple specimens of all *Dryocopus, Celeus*, *Colaptes*, and *Piculus* species to determine whether *D*. *galeatus* plumage traits are novel within *Celeus* and its sister group *Colaptes* + *Piculus*. Given that several of these plumage characters involve subtly different feather tracts that are potentially non-homologous across the Picinae, we chose to map whole phenotype illustrations on the combined data maximum likelihood topology to illustrate the distribution of convergent similarities in appearance among relevant taxa.

## RESULTS

### Sequence attributes

The concatenated sequence alignment contained 3855 characters, of which 1275 were variable and 897 parsimony informative (Table 1). Mitochondrial sequences appeared to be of genuine origin, as stop codons were not observed in open reading frames, base composition was homogenous across samples, and codon specific substitution rates were consistent with known biases. Third position substitutions accounted for 58.7% (527 bp) of the total informative sequence variation, and 69.2% (392 bp) of the informative sites recovered within *Celeus*. By comparison, β-FIBI7 and HMGN2 sequences exhibited little genetic variation, containing just 60 (6.6%) and 85 (9.5%) informative substitutions within the full data matrix respectively, of which 17 (1.9%) and 18 (2.0%) were recovered within *Celeus*. Informative indels were also rare within nuclear sequences; however, a 2 bp deletion shared by all members of *Celeus* and *D. galeatus* corroborate the phylogenetic results presented below (Figure 3). Both specimens of *D*. *galeatus* yielded near identical sequences with no conflict among overlapping amplicons, suggesting an absence of contaminant DNA. All new sequences generated for this investigation have been deposited in GenBank under the accession series KT204492-KT204537.

**Table 1.** Attributes of mitochondrial and nuclear sequence variation in Malarpicini woodpeckers.

| | mtDNA by codon position | | | HMGN2 | β-FIBI7 |
| --- | --- | --- | --- | --- | --- |
|  | 1st | 2nd | 3rd |  |  |
| Length (base pairs) | 744 | 744 | 744 | 709 | 913 |
| Variable Sites (%) | 207 (27.8) | 102 (13.7) | 608 (81.7) | 180 (25.4) | 178 (19.5) |
| Informative Sites (%) | 155 (20.8) | 70 (9.4) | 527 (70.8) | 85 (12.0) | 60 (6.6) |
| Best-fit Model (AIC) | GTR+I+$\Gamma$ | HKY+$\Gamma$ | GTR+I+$\Gamma$ | TVM+ $\Gamma$ | TVM+ $\Gamma$ |
| Frequency %A | 0.3334 | 0.1642 | 0.3420 | 0.2529 | 0.3099 |
| Frequency %C | 0.3218 | 0.3365 | 0.4913 | 0.1658 | 0.1787 |
| Frequency %G | 0.1512 | 0.0951 | 0.0470 | 0.2711 | 0.1782 |
| Frequency %T | 0.1936 | 0.4042 | 0.1197 | 0.3102 | 0.3332 |

**FIGURE 3.**
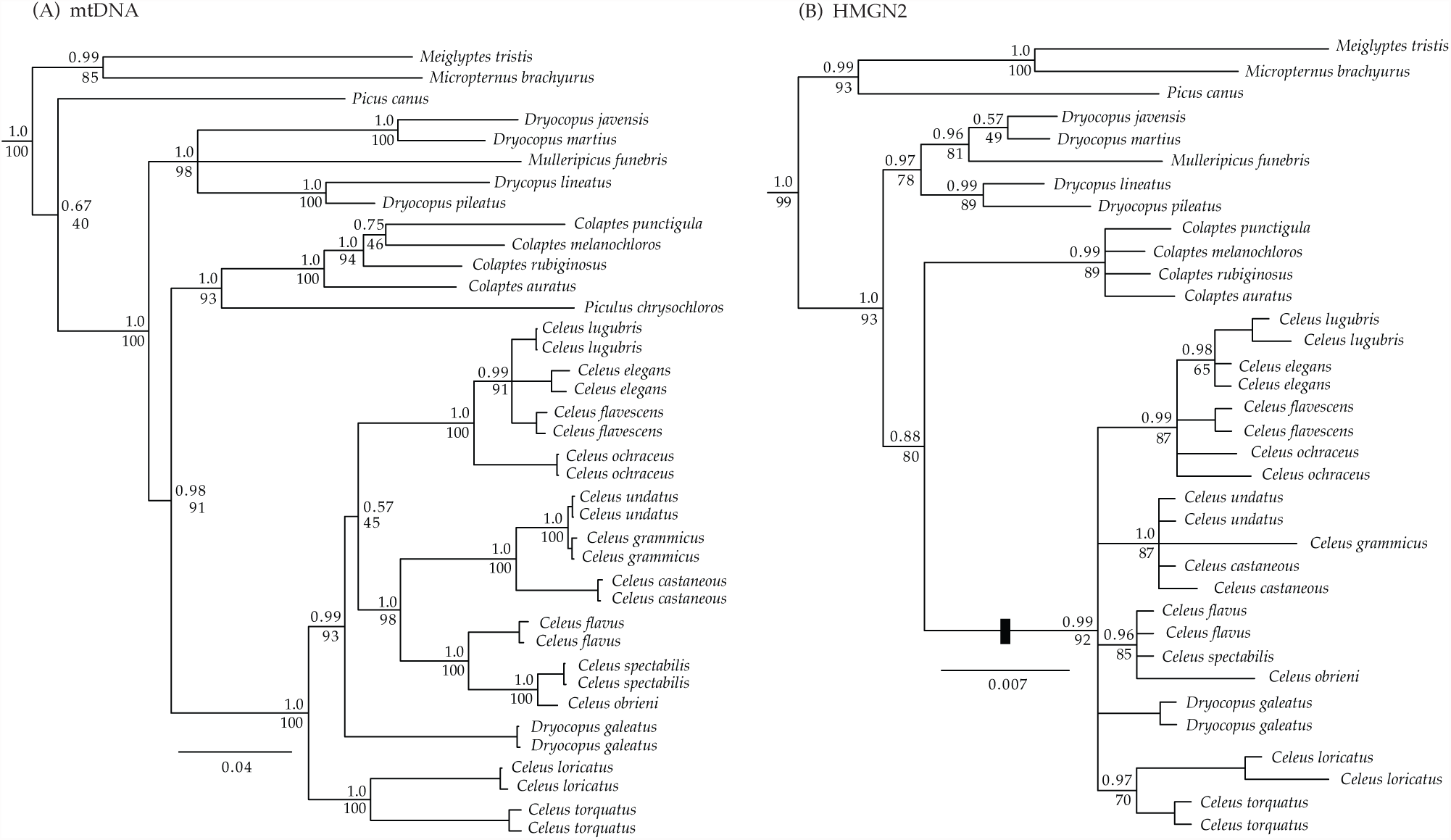
Phylogenetic relationships of the Helmeted Woodpecker *D*. *galeatus* inferred from (**A**) the combined mtDNA data matrix partitioned by codon position, and (**B**) the HMGN2 nuclear locus. Bayesian posterior probabilities and ML bootstrap support values are indicated above and below each node respectively.

### Phylogenetic analysis

Analyses of AIC values generated in jMODELTEST 2 indicated that the general time reversible GTR+I+Г substitution model was most appropriate for the 1^st^ and 3^rd^ codon positions whereas the HKY+Г model was selected for the more conservative 2^nd^ codon position, and a transversion model of evolution (TVM+ Г) was best suited for the HMGN2 and β-FIBI7 nuclear loci (Table 1). Phylogenetic analyses of the individual nuclear loci and the combined four-gene mitochondrial data set recovered similar results, differing primarily in the degree of intrageneric resolution, which reflects the large disparity in rates of evolution and informative variation among these marker sets (Figure 3). As few conflicts were observed among individual gene trees and none were statistically significant, we primarily focus on the combined-data phylogenetic analyses herein, which form the basis of our discussion; however, we do emphasize that both the mitochondrial and HMGN2 analyses strongly reject the monophyly of *Celeus* as currently defined (Figure 3). Phylogenetic results of the β-FIBI7 analyses are provided in Supplemental Material Figure S1, as *D*. *galeatus* was not sequenced for this locus.

Maximum likelihood and Bayesian analyses of the six-gene five-partition data matrix recovered concordant topologies with strong bootstrap and posterior probability support across all but 3 nodes among ingroup taxa (Figure 4). Monophyly of *Celeus* was unequivocally rejected in all analyses, with *D*. *galeatus* placed between the basal *C*. *torquatus* + *C*. *loricatus* lineage (Clade III) and the remainder of *Celeus* diversity. Moderate levels of pairwise sequence divergence recovered between *D*. *galeatus* and other *Celeus* members (8.5 to10.2%; ND2 uncorrected) suggest the former has no close relatives and likely represents an early split within the genus. Although the phylogenetic position of *D*. *galeatus* received strong statistical support in these analyses, weak support between Clade I and II indicate uncertainty in this arrangement, which the nuclear gene tree analyses were also unable to resolve due to a lack of informative variation (Figure 3 and Supplemental Figure S1). This uncertainty had no impact on the position of Clade III, which was consistently recovered as the basal lineage within *Celeus*. The *Colaptes* + *Piculus* clade was placed sister to *Celeus* with moderate to strong support, followed by a paraphyletic *Dryocopus* lineage in which both Old World members of the genus are more closely related to *Mulleripicus* than New World *Dryocopus* (Figure 4). Phylogenetic relationships among outgroup taxa were consistent with previous multi-locus investigations (Benz et al 2006, Fuchs et al. 2007), thus these species were omitted from the final topology to facilitate phenotype mapping within the Malarpicini.

**FIGURE 4.**
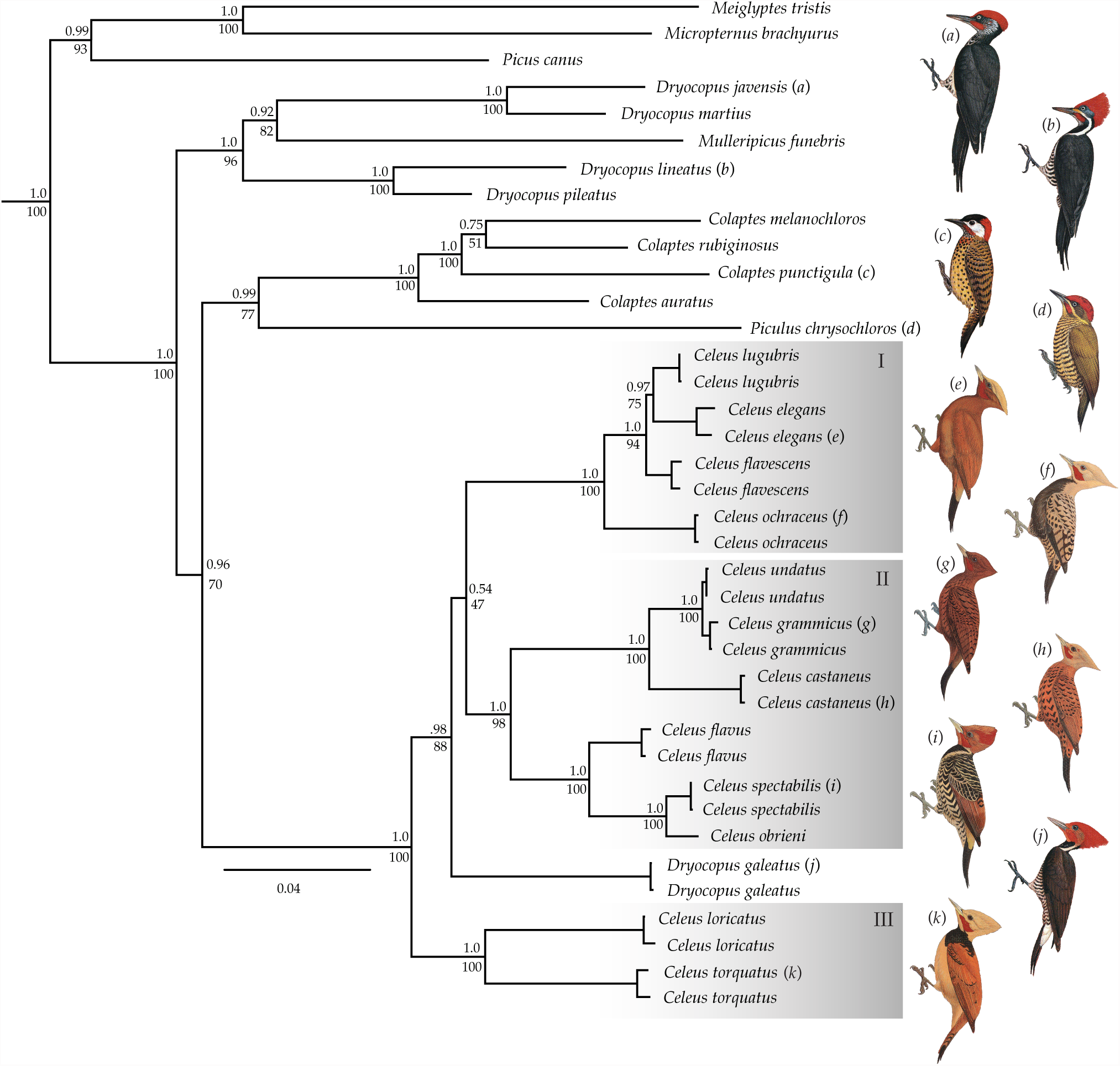
Phylogenetic relationships among Malarpicini woodpeckers inferred from the combined sequence alignment (ND2, ND3, ATP6, ATP8, HMGN2 and β-FIBI7). Bayesian posterior probabilities and ML bootstrap support values are indicated above and below each node respectively. Representative images from each of the three primary *Celeus* Clades illustrate the phenotypic disparity between *D*. *galeatus* and its true congeners.

### Plumage analysis

Similarities in appearance between *D*. *galeatus*, *D*. *lineatus* and *C*. *robustus* are primarily attributed to the fully red crest, pale venter with narrow black banding, and uniform black dorsum (assuming an at rest posture) shared by each of these taxa. Examination of 128 representative study skins encompassing 48 species in 5 genera confirm these traits are absent in all other *Celeus* species, as well as the sister group *Piculus* + *Colaptes*, indicating a convergent evolutionary origin of these traits within *D*. *galeatus* (Figure 4). Although both candidate model species exhibit fine-scale differences in plumage pattern and coloration, *D*. *galeatus* more closely resembles *D*. *lineatus erythrops* in several key aspects. The white lateral neck stripes in *D*. *lineatus erythrops* narrow above the malar and transition to pale orange-brown as they approach the nares, a pattern that is mirrored in *D*. *galeatus* but with little to no stripe definition above the malar and darker cinnamon at the nares (Figure 1). These traits are absent in male *C*. *robustus*, which with the exception of white and black ear coverts have a fully red head, throat, and neck. Although female *C*. *robustus* exhibit a more extensive white and black streak that extends from the ear to the base of the bill, these prominent plumage differences strongly contrast with the head and neck plumage of *D*. *galeatus* and *D*. *lineatus erythrops*. Extensive white scapular patches are characteristic of *D*. *lineatus* populations from northern Mexico to southern Brazil, yet most populations of *D*. *lineatus erythrops* have fully black scapulars, which is also the case in *D*. *galeatus*. Notable phenotypic differences between *D*. *galeatus* and *D*. *lineatus erythrops* include a pale chin with fine black stripping that transitions to an extensive and uniform black upper chest in the later, which clearly contrasts with the cinnamon chin and small black patch restricted to the throat in the former. Nonetheless, these differences are not readily apparent from lateral profiles (Figure 1). Further distinctions include the ivory bill, dark brown irides, and pale rump, which are characteristic of *D*. *galeatus* and contrast with the dark grey maxilla, white irides and black rump in *D*. *lineatus erythrops*.

## DISCUSSION

In his review of avian evolution and homodynamy in morphologically uniform groups, Bock (1963) examined three pairs of woodpecker genera (*Dryocopus*–*Campephilus*, *Gecinulus*– *Blythipicus*, and *Dinopium*–*Chrysocolaptes*) that share highly similar plumage characteristics but exhibit key morphological differences in the foot, tail, and bill; adaptive features that are closely linked to taxon-specific foraging strategies. He concluded that these plumage similarities were evidence of recent shared ancestry, as “the complexity of these color patterns precludes any reasonable possibility of their arising independently in three pairs of genera; no known selection forces could explain such a pattern of convergence”. Over the last decade molecular phylogenetic analyses have confirmed that the inverse is indeed the case, documenting convergent evolution of phenotypic traits in eleven picid genera, and now our data provide yet another example where plumage convergence in the Helmeted Woodpecker has confounded phylogenetic relationships in the Malarpicini (Webb and Moore 2005, Weibel and Moore 2005, Moore et al. 2006, Benz et al. 2006, Fuchs et al. 2008). Although some uncertainty remains with respect to branching patterns among primary *Celeus* clades, our mitochondrial and nuclear data strongly support inclusion of *D*. *galeatus* within the genus (Figures 3 and 4). These results greatly expand *Celeus* phenotypic diversity and demonstrate the independent evolutionary origin of plumage similarities between *D*. *galeatus* and its larger, socially dominant ecological competitors, *D*. *lineatus* and *C*. *robustus*, both of which are syntopic throughout much of the Helmeted Woodpecker’s distribution (Figure 2). By all other accounts, the Helmeted Woodpecker is morphologically highly similar to other *Celeus* species. Its culmen is noticeably curved (similar in shape to that of *C*. *flavescens*), whereas all true *Dryocopus* exhibit distinctly straight and chisel shaped bills that are more robust (wider at the base) and typically darker in coloration. As in all other *Celeus* species, the nares of *D*. *galeatus* are fully exposed and lack the stiff feathering that partially or fully cover the nares of *Dryocopus* species. The dark brown irides of the Helmeted Woodpecker are also characteristic of *Celeus*, whereas all but one species of *Dryocopus* (*D*. *schulzi*) have pale whites irides. An absence of *D*. *galeatus* fluid-preserved anatomical specimens precludes at this time further morphological comparison and diagnosis of internal synapomorphies detailed by Goodge (1972); nonetheless, it appears that previous workers have failed to see the true evolutionary relationships of *D*. *galeatus* largely in part due the historical bias placed on using labile plumage traits to infer phylogenetic relationships, while disregarding other more conservative and potentially informative traits. We suspect that this striking case of plumage convergence likely constitutes a form of interspecific mimicry, whereby *D*. *galeatus* receives social advantages including reduced competitive interference and enhanced access to foraging sites by visually deceiving one or more of its ecological competitors. Although extensive field studies will be required to confirm whether deception is indeed taking place and to what extent ISDM or alternative evolutionary mechanisms are driving these convergent similarities in appearance, the limited behavioral evidence presently available appears to be consistent with the ISDM hypothesis, which we discuss below.

### Non-mimetic evolutionary convergence

Environmental adaptation based on natural selection principles is frequently invoked to explain general phenotypic trends and geographic variation in plumage coloration within birds (Mayr 1963; Zink and Remsen 1986; Hill and McGraw 2006). By contrast, examples of environmental selective pressures driving comprehensive phenotypic convergence of elaborate plumage patterns are relatively uncommon and typically poorly substantiated. Perhaps the best example of the later occurs between *Macronyx croceus* (Motacillidae) of open African savannas, and the meadowlark (Icteridae) species complex *Sturnella spp.*, which inhabit grassland environments throughout much of the New World. The brown streaked dorsal plumage, bright yellow venter, and black pectoral band shared by these allopatric species suggest that there is potential for wholesale phenotypic convergence among taxa exposed to similar environmental selective pressures. Further examples of plumage convergence frequently attributed to environmental adaptation are seen in several oceanic birds including *Alle Alle* and various *Pelacanoides* species, whose black dorsal plumage and white venter are likely related to similar selective pressures associated with foraging in open ocean environments. The proposition that environmental adaptation may be driving the present case of plumage convergence among Atlantic Forest woodpeckers is clearly falsified by the fact that *D*. *lineatus* ranges from northern Mexico to eastern Argentina (Figure 2), occupying a wide diversity of habitats including mangroves, thorn-scrub, open gallery forest, dense rainforest, and pine-oak environments, which collectively range from 0 to 2100 meters above sea level (Short 1982). That any aspect of this species’ visual appearance could be under similar environmental selective pressures in such diverse habitat types is highly unlikely. Moreover, visually similar species in both *Dryocopus* and *Campephilus* further extend the distribution of these phenotypic traits.

Natural and sexual selection processes acting on labile or highly modular phenotypic traits may also foster non-mimetic convergent similarities in appearance (Endler and Théry 1996, Hill and McGraw 2006). Analyses of plumage evolution in New World orioles (*Icterus*) revealed evidence of convergence or reversals in 42 of 44 plumage characters, with repeated evolutionary origins of broad phenotypic similarities in three distinct Clades (Omland and Lanyon 2000). Although several instances of plumage convergence within *Icterus* involve largely allopatric species thereby precluding mimicry or socio-ecological explanations for similarities in appearance, such examples of parallel selection in allopatry appear to be uncommon in birds. Furthermore, examples of convergent plumage evolution among allopatric species typically involve rather coarse plumage details and lack the fine-scale, comprehensive phenotypic similarities necessary for successful visual mimicry. If similar natural or sexual selection processes are responsible for convergent plumage similarities in the Helmeted Woodpecker, why are these traits wholly absent in other *Celeus* species as well as *Colapates* and *Piculus*? Likewise, these hypotheses do not explain why *D*. *galeatus* more closely resembles sympatric populations of *D*. *lineatus erythrops* as apposed to other congenerics or allopatric members of the *lineatus* complex.

### Interspecific visual mimicry

Aposematic forms of mimicry (e.g. Batesian and Müllerian) arise via natural selection processes that typically involve a third-party observer in addition to the model and mimetic taxa (Ruxton 2004). In the absence of aposematic signaling for predation avoidance, interspecific visual mimicry is thought to confer social advantages to species in close ecological competition by enabling mimetic taxa to avoid aggression through deception and increase their foraging opportunities (Wallace 1869, Diamond 1982, Rainey and Grether 2007, Prum and Samuleson 2012). Diamond (1982) hypothesized that competitive mimicry reduces the frequency of attack by larger model species, thereby altering the social hierarchy at highly contested foraging sites counter to size mediated social dynamics; however, he remained skeptical of whether selection pressures associated with the risks of aggression were sufficient to prevent models attacking the smaller mimetic species. As such, Diamond did not rule out the possible role of a third party observer in promoting and maintaining competitive mimicry between *Philemon* and *Oriolus* species.

More recently, Prum and Samuleson (2012) used evolutionary game theory modeling to confirm that intraspecific attacks are indeed constrained by the costs of aggression, thereby fostering social opportunities for interspecific competitive mimicry purely within the context of a two party system. Fitness dynamics derived from these game theory models predict the evolution of ISDM when associated costs of mimicry are low, the background fitness of the mimetic species is greater than that of the model, and the values of contested resources are neither exceedingly high nor low. The ISDM hypothesis also predicts that selection on the dominant model species will favor evolution of divergent phenotypic traits to reduce the efficacy of mimicry and its associated costs. In time, these counter selective forces may lead to co-evolutionary radiations between model and mimetic taxa as seen in the *Philemon* and *Oriolus* example or perhaps between *Dryocopus* and *Campephilus* (Diamond 1982, Prum 2014). Lastly, size differences between mimic and model are constrained in ISDM such that visual deception must be feasible at distances relevant to their behavioral ecology. The later prediction is readily testable and should provide critical insight into the functional significance of phenotypic convergence in birds, as environmental based non-mimetic hypotheses do not predict explicit size relationships between model and mimic, nor do previous mimicry hypotheses of Moynihan (1968) and Cody (1969).

In his recent review of avian visual mimicry, Prum (2014) proposed fifty phylogenetically independent examples of ISDM from thirty families. The average mimic body mass was 55.7% of the putative model counterpart and a linear regression of body mass between subordinate mimic and dominate model revealed a strong positive correlation with a slope of 0.5684 and *R*^*2*^ value of 0.83. This close association between asymmetry in body size and similarity in appearance strongly suggest these convergent signals are evolving in the context of competitive mimicry to facilitate interspecific deception of both the mimic’s identity and body size. By simply appearing to be large rather than physically evolving larger body size to legitimately dominate an ecological competitor, the mimetic species may incur advantages in physiological efficiencies while simultaneously having access to a wider diversity of ecological resources, translating to greater adaptability and evolutionary persistence. The selective pressures for such advantages may be especially acute in groups that exhibit highly specialized foraging strategies, which may explain the high prevalence of phenotypic convergence in the Picinae.

### Visual mimicry in the Helmeted Woodpecker

Interspecific mimicry in *D*. *galeatus* was first proposed by Willis (1989), who identified the larger Robust Woodpecker (*Campephilus robustus*) as a possible dominant model (Figure 1), given that both occasionally forage together in mixed-species flocks. In light of our phylogenetic results, it’s plausible that *galeatus* is a mimic of both *C*. *robustus* and *D*. *lineatus*; however, the later bears greater similarity to *galeatus* given its conspicuous white neck stripes and darker lores. Moreover, we underscore the fact that *galeatus* is sympatric with southern populations of *D*. *lineatus erythrops*, both of which lack the white scapular patches that are otherwise characteristic of the broadly distributed *lineatus* complex (Winkler and Christie 2002). At approximately 28 cm in length, the Helmeted Woodpecker is 22 to 25% smaller than *D*. *lineatus erythrops* (36 cm) or *C*. *robustus* (37 cm) respectively, but weighs less than half of either model taxon. This disparity in size and weight undoubtedly confers a substantial physical advantage to the larger *Dryocopus* and *Campephilus* models, as even small differences (< 10%) in body mass can lead to greater success in competitive interference for numerous avian groups (Ford 1979, Mauer 1984, Milikan et al. 1985; Atalo and Moreno 1987, Robinson and Terbourgh 1995). Critically, the size differences observed between galeatus and either model species appear to be consistent with the ecological and psychophysical constraints required for visual deception as outlined by Prum (2014). Given that the difference in distance between two objects sharing the same visual angle scales linearly with difference in size, mistaking a mimic species for conspecific dominant model would require overestimating the true distance by just 32 %, which is well within the ecological context these species regularly encounter one another. Although avian visual acuity varies substantially among taxonomic groups, ophthalmological research and psychophysical data suggest visual deception of this nature for non-raptorial birds is possible at a distance of three meters or more (Hodos 1993, Prum 2014).

Knowledge of the Helmeted Woodpecker’s ecology and foraging behavior remains extremely limited in comparison with other Neotropical picids, however *galeatus* appears to be an ant specialist (*Crematogaster* sp.) that regularly consumes small fruits such as *Alchornea sidifolia* berries (Santos 2008, Lammertink et al. 2012, and Zimmer pers. obs.). This secretive species primarily forages at mid-levels on interior branches, quietly probing rotting wood, which is consistent with *Celeus* foraging behavior (Short 1982, Winkler and Christie, 2002). All twelve species currently recognized within *Celeus* are documented ant or termite specialists that regularly consume fruits and rarely exhibit strong excavating or bark scaling behavior characteristic of *Dryocopus* and *Campephilus*. Both *C*. *robustus* and *D*. *lineatus* consume fruits and the later taxon regularly forages on ants including *Crematogaster*, *Azteca*, and *Camponotus* species. The diet of these larger picids differs from that of *galeatus* in that both *Dryocopus* and *Campephilus* species use their powerful bills to excavate beetles and their larvae from deep within rotten to semi-rotten substrates. Given that rotting trees of appropriate age and decomposition are often in limited supply within forest environments, competition for suitable foraging substrates rather than direct competition for a particular food species may be responsible for the evolution of competitive mimicry between *galeatus* and either dominant model species. Although southern populations of *D*. *lineatus erythrops* appear to forage within higher strata than either *galeatus* or *C*. *robustus*, the competition for foraging sites may encompasses an entire tree given that most *Dryocopus* and *Campephilus* species generally do not tolerate unfamiliar conspecifics in the vicinity of an active feeding site. Despite an absence of detailed knowledge on the socio-ecological interactions between this trio of woodpecker taxa, evidence of interspecific mimicry is most consistent with the ISDM hypothesis presented by Prum and Samuelson (2012), as woodpeckers are not known to sequester toxins in their skin or feathers and previous suggestions by Cody (1969) that phenotypic convergence promotes enhanced interspecific territoriality is disproven by the fact that both *galeatus* and *C*. *robustus* occasionally attend mixed species flocks (Willis 1989). Moreover, Cody’s hypothesis does not take into account the strong asymmetry in body size associated with cases of competitive mimicry, casting doubt on the evolutionary stability of mutual exclusion. Moynihan’s (1968) hypothesis is not applicable in the present case as neither mimic nor model species are facultative mixed-flock attendants. Likewise, little support for this hypothesis is seen in other proposed examples of avian competitive mimicry (Diamond 1982, Prum 2014).

Behavioral investigations examining the socio-ecological circumstances associated with cases of avian phenotypic convergence will be required to confirm the prevalence of ISDM versus traditional three party mechanisms of visual mimicry. Although Prum and Samuleson (2012) demonstrate that the evolution and persistence of competitive mimicry is possible exclusively within a two-party system, it seems plausible that ISDM may operate synergistically with third-party deception mechanisms, as the potential pool of non-model ecological competitors is much larger. Given that most birds communicate a broad array of intra and interspecific information through vocalizations, future field studies examining potential cases of competitive mimicry must also take into account the mimic’s vocal behavior particularly in the presence of models and other heterospecific competitors, as any discontinuity between morphological and behavioral mimicry would likely preclude the possibility of deception. As such, we predict mimetic taxa will vocalize less in the presence of ecological competitors by comparison with their non-mimetic congeners. Although little is known of the Helmeted Woodpecker’s vocal behavior, it appears to call less frequently than other members of *Celeus*, which may account for the dearth of visual sightings and possibility of extinction reported by Short (1982).

### Conservation status and taxonomic implications

The Helmeted Woodpecker inhabits semi-deciduous and mixed-forest environments from São Paulo, Paraná, and Santa Catarina in southeastern Brazil, west to eastern Paraguay, and south to Misiones in extreme northeastern Argentina (Short 1982, Collar et al. 1992, Hayes, 1995, Winkler and Christie 2002). This little-known species has undergone dramatic population declines and vanished from much of its former distribution in the later half of the 20^th^ century due to extensive regional deforestation (Short 1982, Galindo-Leal and de Gusmão Câmera 2003, Santos 2008). Consequently, it is currently listed as Vulnerable by BirdLife International and considered to be among the rarest of Neotropical woodpeckers (Lammertink et al. 2012, BirdLife International 2013). Although new details of its ecology and natural history are slowly emerging, the conservation status of *D*. *galeatus* remains unclear and deserves careful examination given regional trends in habitat loss.

In light of our phylogenetic data, described morphological differences (Short 1982), distinct vocalizations (*galeatus* has similar vocalizations to *Celeus torquatus* and *C*. *flavus*; XC61093, Xeno-Canto, http://www.xeno-canto.org), and mechanical sound production (drumming of *galeatus* is *Celeus*-like; XC24502, XC17067, Xeno-Canto) the Helmeted Woodpecker clearly requires reclassification as *Celeus galeatus*.

## ACKNOWLEDGMENTS

We thank the following institutions and their staff for providing tissue loans: American Museum of Natural History (AMNH), Field Museum of Natural History (FMNH), National Museum of Natural History (USNM), Louisiana State University Museum of Natural Science (LSUMNS), Museo de Zoologia, Facultad de Ciencias, Universidad Nacional Autonoma de Mexico (UNAM), University of Kansas Biodiversity Institute (KU), University of Washington Burke Museum (UWBM), and Los Angeles County Museum of Natural History (LACM). We are also grateful to the many field collectors that made this research possible. This work was supported by the Monell Molecular Laboratory and Cullman Research Facility in the Department of Ornithology, American Museum of Natural History.

## Appendix A. Taxon sampling used in this study.

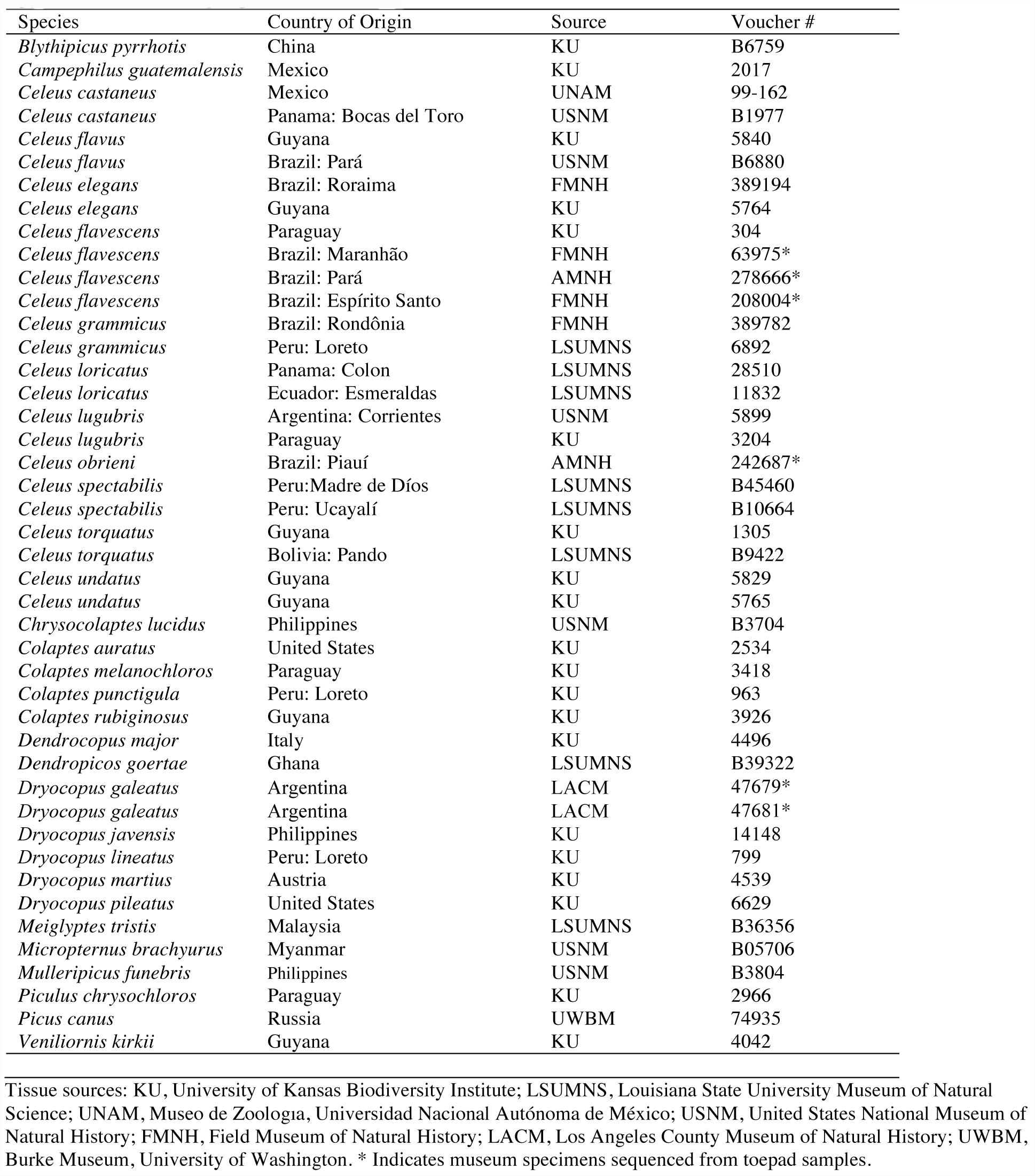

**Supplemental Figure S1.**
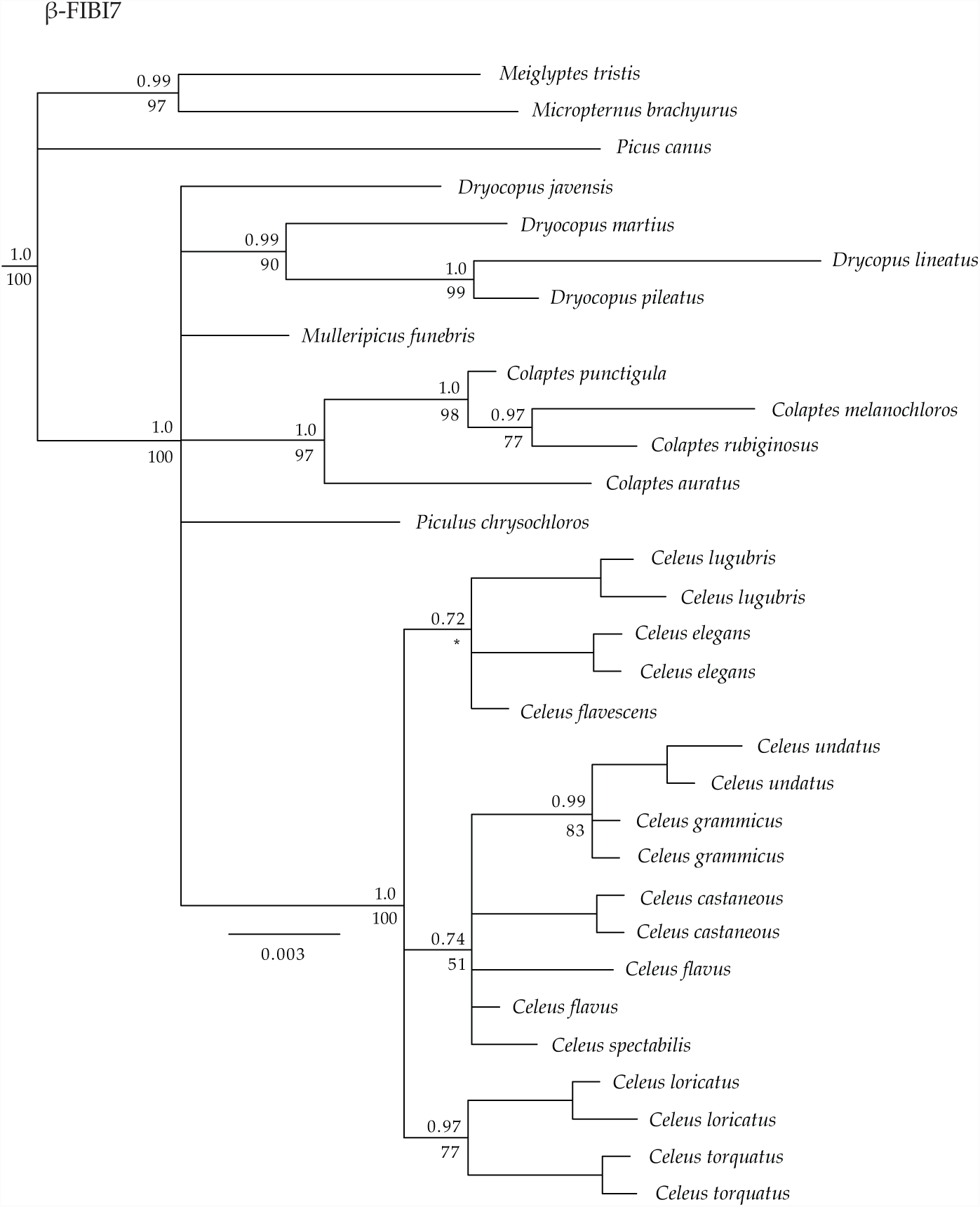
Phylogenetic relationships among the Malarpicini inferred from β-FIBI7 sequence data. Bayesian posterior probabilities and ML bootstrap support values are indicated above and below each node respectively.

